# Mechanisms of Cholesterol Binding to LAT1

**DOI:** 10.1101/2024.01.02.573920

**Authors:** Keino Hutchinson, Avner Schlessinger

## Abstract

The human L-type amino acid transporter 1 (LAT1; SLC7A5), is an amino acid exchanger protein, primarily found in the blood-brain-barrier, placenta, and testis, where it plays a key role in amino acid homeostasis. Cholesterol is an essential lipid that has been highlighted to play a role in regulating the activity of membrane transporters such as LAT1, yet little is known about the molecular mechanisms driving this phenomenon. Here we perform a comprehensive computational analysis to investigate cholesterol’s role in LAT1 structure and function, focusing on four cholesterol binding sites (CHOL1-4) identified in a recent LAT1-apo inward-open conformation cryo-EM structure. We performed four independent molecular dynamics (MD) simulations of LAT1 bound to each cholesterol molecule, as well as molecular docking, free energy calculation by MM/GBSA, and other analysis tools, to investigate LAT1-cholesterol interactions. Our results indicate that CHOL3 provides the most stable binding interactions with LAT1, and CHOL3 and CHOL1 sites have the largest stabilizing effect on LAT1’s primary functional motifs (hash and bundle) and substrate binding site. Our analysis also uncovers an alternative cholesterol binding site to the originally assigned CHOL1. Our study improves the understanding of cholesterol’s modulatory effect on LAT1 and proposes candidate sites for discovery of future allosteric ligands with rational design.

## INTRODUCTION

In humans, amino acids are an essential many physiological and cellular processes including protein formation, signaling molecules, and energy homeostasis^1^. Membrane transporters play a key role in mediating the movement of amino acids across biological membranes in cells and organs^2^. The L-type amino acid transporter 1 (LAT1; SLC7A5) imports large neutral amino acids such as phenylalanine and tyrosine^3^, in a 1:1 exchange for intracellular amino acids (e.g., histidine)^4^. LAT1 is a member of the heteromeric amino acid transporters (HATs)^5^, which are typically composed of a heavy and a light chain. HATs are typically comprised of two families of proteins: SLC3 (heavy chain) and SLC7 (light chain), which are linked by a conserved disulfide bridge^6^.

Cholesterol is an important sterol^7^ that plays a key role in maintaining membrane integrity and can directly interact with integral membrane proteins^8^, including membrane transporters^9^ and G-protein coupled receptors (GPCRs)^10^. Atomic resolution structures of membrane transporters have shown how cholesterol interacts with these proteins^11–13^, and multiple recent studies showed that this interaction can modulate transport activity^14–18^. For example, the depletion of membrane cholesterol with the cholesterol chelating agent methyl-β-cyclodextrin (MβCD) reduced the affinity (K_m_) and uptake velocity (V_max_) for substrate uptake through the dopamine transporter (DAT)^14^, a distant homolog of LAT1. Comparatively, increasing cholesterol levels in the membrane, promotes DAT to adopt an outward-open conformation^15^. Similarly, for the serotonin transporter (SERT), cholesterol depletion with either MβCD, cholesterol oxidase or cholesterol-binding fluorochrome filipin, results in the decrease of SERT activity in the form of lowered K_m_ of substrate and ligand binding as well as the V_max_ of transport^16^. Taken together, these studies suggest that cholesterol can play a central role in stabilizing transporters in a specific state (e.g., outward or inward open) of their transport cycle and affect substrate interaction. Recent studies have suggested that cholesterol has a similar modulatory effect on LAT1 to that seen in its homologs SERT and DAT^19, 20^. Using a LAT1 mediated L-DOPA uptake assay, Dickens et al. showed that the depletion of cellular cholesterol by MβCD, reduced the V_max_ but not the K_m_ of transport. Similarly, Cosco et al, proposed that cholesterol promotes a stabilization of the inward open conformation of LAT1 and that there may also be a synergistic effect with ATP^20^. Interestingly, using computational methods, they identified a putative cholesterol binding site that is potentially important for modulating ATP binding.

Molecular Dynamics (MD) simulations is a useful tool for examining protein-lipid interactions^21^. Atomistic and Coarse-grained (CG) MD allows for the analysis of specific direct interactions between lipids, mainly cholesterol, and provide insight on membrane protein stability^22–24^. With no assigned lipid, CG force fields, such as MARTINI^25^, can be used in an unbiased manner to investigate potential lipid binding sites with sufficient sampling due to the decreased degrees of freedom of the underlying model^26–29^. Additionally, with an increasing number of membrane proteins being solved at near atomic resolution with assigned lipids^30, 31^, MD simulations can be used to characterize this lipid interaction and its effect on modulating protein function, and particularly transporters^18, 32^.

In this study, we perform a comprehensive computational analysis of LAT1-cholesterol binding sites (CHOL1-4) in LAT1 structure^33^. Specifically, we first map various structural features of each CHOL, including the residue conservation, and binding pocket properties. We then perform MD simulations to assess specific interactions between LAT1 and each cholesterol molecule and further analyzed the effect on LAT1’s primary functional motifs (hash and bundle) and substrate binding site. We also report an alternative CHOL1 binding pocket than the assigned pocket in the cryo-EM structure, which was revealed during the CHOL1 MD simulation. Next, we assess which of the sites are amendable to a cholesterol docking by performing molecular docking and MM/GBSA calculation. Finally, we discuss the role of cholesterol binding sites may have on regulating LAT1 activity, as well as their relevance for drug discovery.

## MATERIALS AND METHODS

### Protein preparation

The coordinates of LAT1 (chain B) were extracted from its cryo-EM structure in the apo inward-open conformation (PDB: 6JMQ)^33^, including four cholesterol molecules bound to chain B. The cryo-EM structure contained two unstructured loops at residues 220-230 and 353-369 were modeled with MODELLER^34^, using two cryo-EM structures of LAT1 in the inward occluded conformation (PDB: 6IRT^35^, 6IRS^35^), as templates.

### Residue conservation

A multiple sequence alignment (MSA) of the human SLC7 family members was generated using MUSCLE^36^ via the EMBL-EBI search and sequence analysis tools^37^ (Suppl. Fig. 1). The resulting MSA was used as input into the ConSurf, a server that estimates and visualizes the per-residue conservation of proteins^38^. ConSurf calculates a normalized score, which is then placed on scale from 1-9, which scores the relative degree of conservation for each amino acid position, where 1 denotes highly variable and 9 conveys highly conserved. This score is then mapped onto the protein structure for easier visualization (Fig. 1A) of regions of high or low conservation and visualized in PyMOL^39^. The average conservation of each site (sum of per-residue conservation score / total number of residues) was defined by residues within 5Å of each assigned cholesterol.

**Figure 1:**
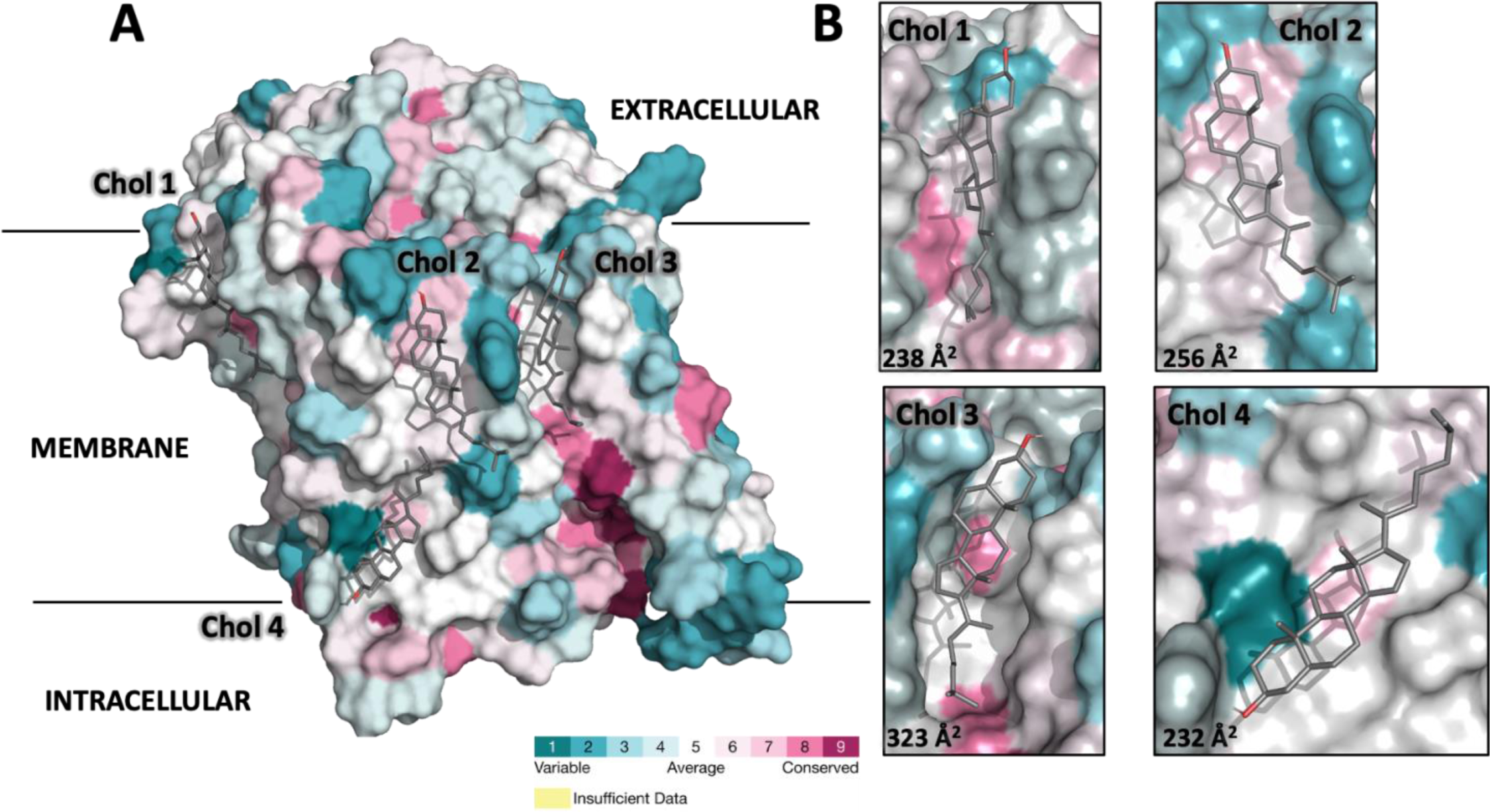
Structural features of the assigned cholesterol binding sites in LAT1. (**A**) Surface representation of LAT1 cryo-EM structure in the apo inward open conformation (PDB:6JMQ). Residue conservation among all members of the human SLC7 family generated using ConSurf (Methods). The colors correspond to the degree of conservation, where red, white, and cyan indicate conserved, average, and variable residues, respectively. The average per-residue conservation of each site was calculated to be: 4.4, 3.5, 4.1 and 3.7 for CHOL1-4 respectively. (**B**) Zoomed in view of the cholesterol binding sites (CHOL1-4) with the corresponding surface area of each site in Å^2^ calculated using POVME. All visualizations were generated using PyMOL.

### Pocket volume measurement

POVME3^40^ (Pocket Volume Measurer 3) was used to calculate binding site volume. We used default parameters for Ligand-defined inclusion region, using the four assigned cholesterol molecules from the inward-open structure (PDB:6JMQ), as the reference ligands.

### Cholesterol binding pocket features

The descriptors of the binding pocket of CHOL1-4 were calculated using dpocket, a pocket descriptor value generator, from the fpocket^41^ open source platform. We defined each site using the coordinates of cholesterol in each CHOL binding pocket. The following features were assessed using dpocket: pocket volume, pocket solvent accessible surface area, pocket polar and apolar surface area, number of alpha spheres, alpha sphere density, mean alpha sphere solvent accessibility, proportion of apolar alpha spheres, hydrophobicity score, volume score, polarity score, charge score, proportion of polar atoms, and maximum alpha sphere distance.

### MD simulations

We used the model generated after loop modeling (see above), as input protein structure for the molecular dynamics (MD) simulations. The four assigned cholesterols from the cryo-EM structure were also separated into individual protein + lipid models (CHOL1-4). The starting membrane bi-layer model was prepared using CHARMM-GUI^42^. 452 POPC lipids were added to the system along with 7,636 water molecules in a triclinic periodic box with 131×131×102nm dimensions. 81 Na^+^ and 81 Cl^-^ ions were added to ensure charge neutrality and a salt concentration of 0.15 M. The CHARMM36m^43^ force field was used for the protein, lipids, and ions; and the TIP3P model were used for the water molecules, respectively.

All MD simulations were performed using the GROMACS^44^ simulation package version 2020.3. A 100 nanosecond-long equilibration was performed using GROMACS following the standard CHARMM-GUI protocol, consisting of multiple consecutive simulations in the NVT and NPT ensembles. During these equilibration steps, harmonic restraints on the positions of the lipids, ligand, and protein atoms were gradually switched off. Each production MD simulation was simulated for 200ns, with the output coordinates of the simulation saved every 100 ps. The temperature of the system was kept constant at 310K, with the Nosé-Hoover scheme and pressure constant of 1 atm using a Parrinello-Rahman approach with semi-isotropic scaling.

### MD simulation trajectory analysis

Root mean square deviation (RMSD) analysis was calculated using the fast QCP algorithm in MDAnalysis^45^. The initial state of the system was defined as first frame of the production trajectory and compared to the corresponding time point of the simulation.

The interaction map for each cholesterol at CHOL1-4 was generated using ProLIF^46^, with the trajectory of each individual MD simulation used as input. The following interactions were assessed to generate the interaction maps for CHOL1-4: ‘Hydrophobic’, ‘HBDonor’, ‘HBAcceptor’, ‘PiStacking’, ‘Anionic’, ‘Cationic’, ‘CationPi’, ‘PiCation’, ‘VdWContact’. Each specific interaction is determined once the ligand (cholesterol) and protein (LAT1) residues satisfy geometrical constraints based on typical distances (Å) and/or angles (deg)^46^. The generated interactions are then tabulated for analysis.

The fluctuation of the binding site volume was obtained with mdpocket^47^ from the trajectory of each independent CHOL simulation.

Clustering of LAT1 and cholesterol binding poses were performed in two steps using: i) RMSD calculated on all the heavy atoms of the ligand and of the protein residues within 5 Å of the ligand cholesterol and ii) the gromos^48^ algorithm with a cutoff equal to 1.5 Å.

### Molecular docking

Molecular Docking was performed using Glide^49^ from the Schrödinger suite. Cholesterol was docked to CHOL1-4, obtained from the cryo-EM structure of LAT1 in the apo inward-open conformation (PDB: 6JMQ). These structures were prepared for docking with the Glide Protein Preparation under default parameters. The ligand binding site was defined based on the coordinates of the respective ligand in each published structure, except for CHOL1^refined^, which came from cluster 1 of the CHOL1 simulation. The receptor grid for docking was generated via Glide Receptor Grid Generation panel. The cholesterol used in molecular docking were prepared using LigPrep of the Schrödinger suite using default parameters. The docking results were visualized via PyMOL.

### Free energy calculation by MM/GBSA

Molecular Mechanics with generalized Born and surface area solvation (MM/GBSA) was calculated using the Prime MM/GBSA^50^ calculator from the Schrödinger suite. The binding pose generated from molecular docking was used as an initial input for MM/GBSA calculation.

### Data availability

Scripts used for post md simulation trajectory analysis are available at: https://github.com/schlessinger-lab/LAT1-cholesterol

## RESULTS

### Description of putative cholesterol binding sites

In our initial analysis, we characterized the structural attributes of four putative cholesterol binding sites of LAT1. Our focus was on residue conservation, electrostatic potential, as well as other molecular features (e.g., hydrophobicity and volume) of CHOL1-4 (Methods). Our findings indicated that while these sites did not display high global residue conservation, localized regions within them exhibited significant conservation, achieving scores of 6 or higher (Fig. 1A). Interestingly, on average, the residues in CHOL1 and CHOL3 are more conserved than those in CHOL2 and CHOL4 among the SLC7 family, with CHOL3 presenting two distinct patches of high residue conservation (Fig. 1B).

Further analysis of the electrostatic potential revealed a dichotomy between the sites: CHOL1 and CHOL4 were characterized by neutral to hydrophobic charges, while CHOL2 and CHOL3 had pockets of negative charge amidst an otherwise hydrophobic landscape (Suppl. Fig 2). This pattern aligns with established literature indicating that cholesterol binding sites in membrane proteins^51^ generally exhibit a hydrophobic character interspersed with minor polar regions, often comprising of Asn, Gln, and/or Tyr residues that typically interact with cholesterol’s β-hydroxyl group^51^.

Expanding our investigation to molecular descriptors using the dpocket scoring function^41^ (Methods) and the surface area and volume of each site. Here, the binding pockets varied notably in size. For instance, CHOL3 not only possessed the largest surface area at 323 Å^2^ (26% larger than the second best), but also ranked as the most hydrophobic alongside CHOL4. Additionally, CHOL3 exhibited the highest alpha sphere density and the most substantial convex hull volume, indicative of the cavity’s size and depth (Suppl. Table 1). When juxtaposing CHOL1-4 against known cholesterol or lipid densities reported in LAT1 structures^19, 20, 35, 52^, CHOL3 corresponded with a distinct “cholesterol-like density” observed in the inward-occluded cryo-EM structure complexed with the LAT1 inhibitor BCH. Taken together, these results suggests that CHOL3 is potentially the most conducive site for robust cholesterol interaction.

### MD Simulations of cholesterol in CHOL1-4

Next, we investigated the stability of cholesterol-LAT1 interactions in each site by performing MD simulations. The CHOL3 simulations (Fig. 2) displayed that cholesterol becomes relatively stable at around 4Å from the initial pose, after exhibiting an initial fluctuation (i.e. after 60ns), which is mainly due to cholesterol binding deeper into the binding pocket from the initial binding pose (Suppl. Mov. 1 and Suppl. Fig 3. A). This movement is captured by clustering analysis of the cholesterol’s movement during the simulation (Suppl. Fig 3C). In addition, the cholesterol in CHOL1 exhibited a similar trend of initial fluctuation followed by a stable binding pose after around 75ns (Fig. 2). Notably, the cholesterol in CHOL1 moves lower down the protein (closer to the intracellular membrane), than the initial placement observed in the experimentally determined structure (Suppl. Fig 3A). Finally, the cholesterols in CHOL2 and CHOL4 do not stabilize at either binding site, with CHOL4 being highly unstable and eventually moving away from the protein (Fig. 2). These results provide further support that cholesterol has the most complementarity with CHOL3.

**Figure 2:**
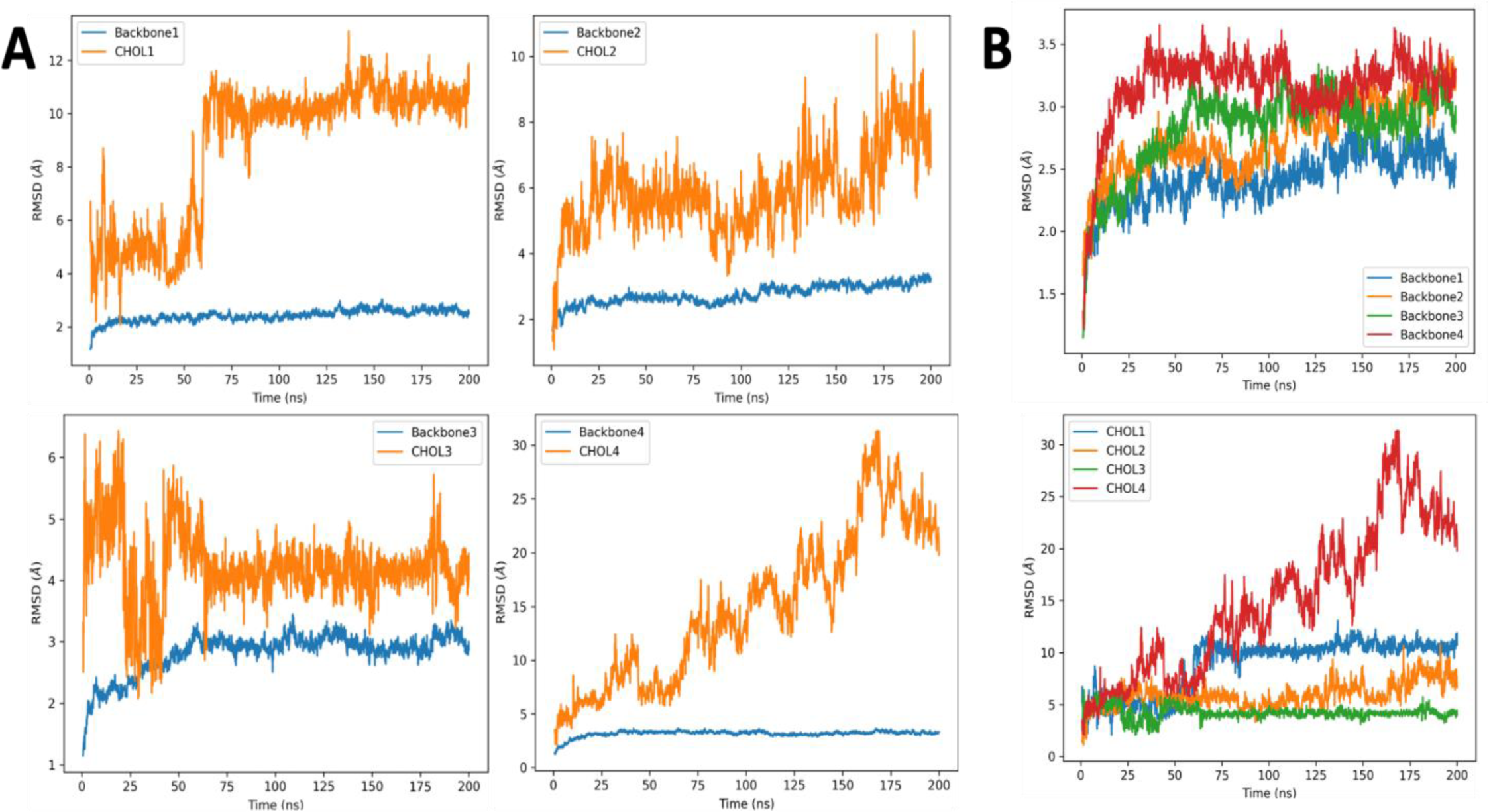
Quantitative analysis of stability of assigned cholesterols and protein backbone. **(A)** Root Mean Square Deviation (RMSD) analysis of four independent MD simulations focusing on the movement of the backbone α-carbon atoms of the inward open conformation and an individual cholesterol at each binding site (CHOL1-4). The x-axis displays the duration of the simulation trajectory (in nanoseconds), and the y-axis shows the movement of atoms in angstroms (Å), compared to the initial frame of the trajectory. **(B)** Comparison of the backbone atoms (above) of all four CHOL simulations and the cholesterols at each site (below), from the four simulations shown in (A).

### Molecular interactions between cholesterol and CHOL1-4

We then evaluated what interactions, if any, mediate cholesterol binding in each site by generating ligand interaction maps from each MD Simulation (Methods). Notably, cholesterols at CHOL3 and CHOL1 made the highest total number of contacts with unique residues (11 each). Furthermore, CHOL3 and CHOL1 had multiple residues make contacts with cholesterol for over 90% of the simulation (Fig. 3). In contrast, we observed no interactions at CHOL2 and CHOL4 that were sustained for over 85% of the simulation and CHOL4 had a single interaction for approximately 55% of the simulation.

**Figure 3:**
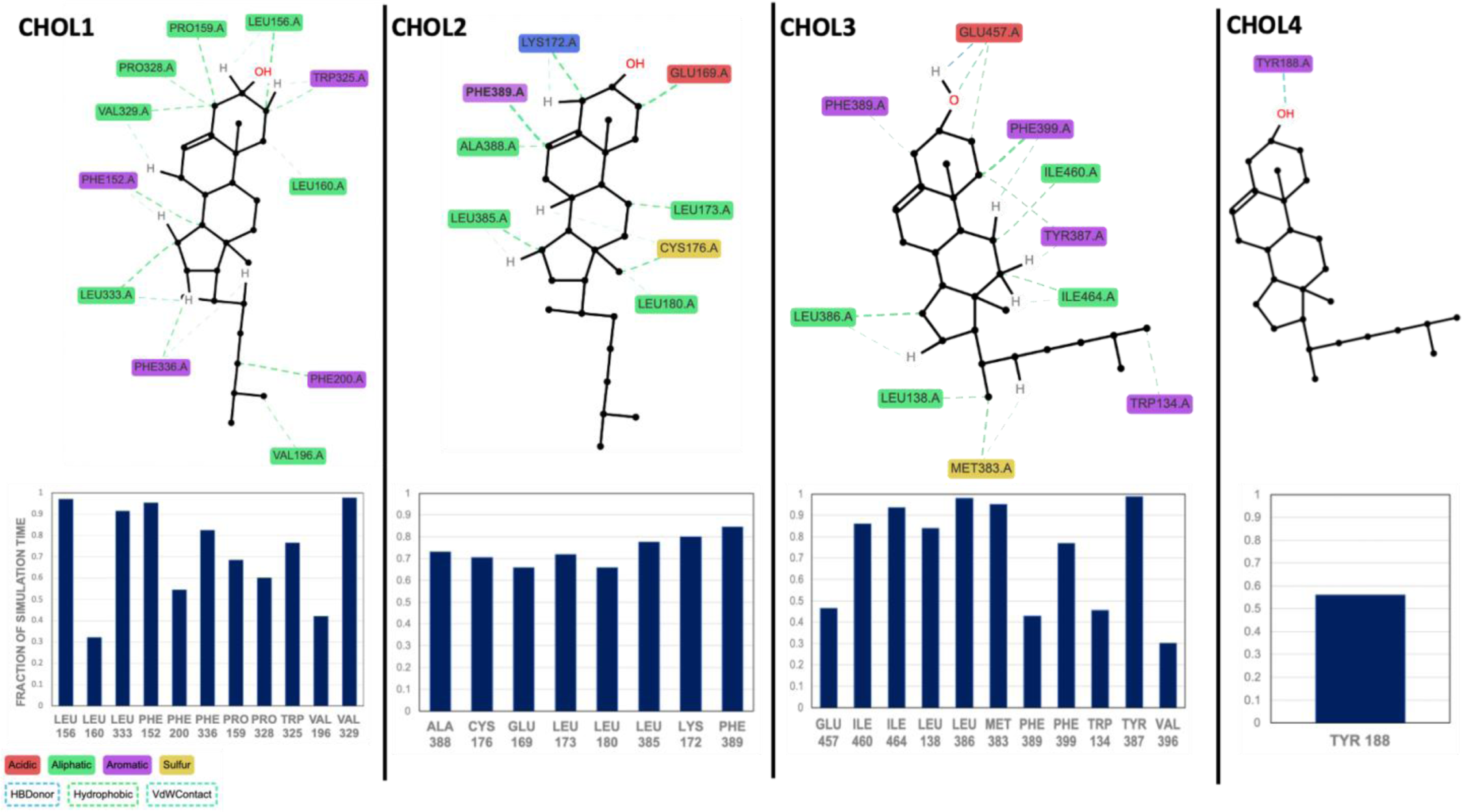
CHOL1-4 interaction map. Specific interactions between cholesterol and residues at each putative cholesterol binding site (CHOL1-4) are shown with dashed lines, including hydrogen bonds (blue), hydrophobic interactions (green), and Van der Waals contacts (green). Amino acids are colored based on their sidechain properties, including acidic (red), aliphatic (green), aromatic (purple), and sulfur (yellow). All interactions are derived from each individual MD simulation (CHOL1-4). The fraction of time during the individual simulations that each residue formed interactions with cholesterol is summarized in a table (below).

We then examined which elements of cholesterol make interactions at each CHOL. In brief, we divided cholesterol into 3 key moieties: (i) head (hydroxyl group), (ii) core (tetracyclic rings (A–D) system), and (iii) tail (flexible iso-octyl group). For CHOL3, we observe that the head group made a hydrogen bond with E457 of LAT1 which was maintained for around half of the simulation time (Figure 3). Interestingly, this interaction is not observed in the native pdb structure but is formed only during the simulation. Additionally, both the core (C1-F399, C12-I464, C15-L386) and tail (C21-L138, C26-W134) moieties of cholesterol, form multiple hydrophobic (van der Waals) interactions with LAT1. In CHOL1, the cholesterol’s core (C2-L156, C3-V329, C14-F152, C15-L333) and tail (C24-F200, C26-V198) make hydrophobic interactions with LAT1, but the head does not make any contacts. As expected, these hydrophobic interactions occurred with residues further away from the extracellular space, matching the cholesterol rearrangement observed in Fig. 2. Comparatively, cholesterol at CHOL2 only interacts via the core (C2-E169, C4-K172, C6-F389, C11-L173, C18-C176, C15-L385) and not the head or tail. The lack of the contacts at theses moieties, provide a rational explanation for the increased flexibility and lack of stabilized binding pose observed in Fig. 2. In CHOL4, cholesterol only forms a transient interaction with LAT1 during the simulation, through its head group (i.e., a hydrogen bond with Y188), rationalizing why cholesterol binding is not maintained at the site.

Taken together, these results suggest that residues in CHOL3 and CHOL1 interact with multiple key elements of cholesterol, which aids in stabilizing cholesterol binding to LAT1. CHOL2 is amenable for cholesterol binding but this may be more transient in nature due to the flexibility of the tail and lack of hydrogen bonding. Lastly, CHOL4 is highly shallow and lacks specific interactions with LAT1.

### Cholesterol’s effect on LAT1 functional motifs and substrate site

To evaluate the functional relevance of cholesterol binding to CHOL1 and CHOL3, we investigated if the binding of cholesterol at these sites is correlated with structural differences in LAT1’s functional motifs and orthosteric substrate binding site. In brief, LAT1 has 12 Transmembrane helices (TMs), with 2 functional motifs: the “rocking” bundle (TMs 1, 2, 6, 7) and the hash (TMs 3, 4, 8, 9), which are thought to mediate conformational transition and substrate binding^33, 35, 52^ (Fig. 4A). The inward-open conformation is structurally defined by the rocking bundle (mainly TM1a and TM6b) being rotated 27° away from the hash domain, when compared to the outward occluded state^52^, which exposes the substrate site to the intracellular side of the cell. Thus, we compared the stability of the backbone α-carbons of these two functional motifs, along with residues in the substrate binding site. Our results suggest that cholesterol binding at CHOL3 stabilizes both domains and the substrate binding site, which all appear to move in a coordinated fashion after 50ns (Fig. 4B).

**Figure 4:**
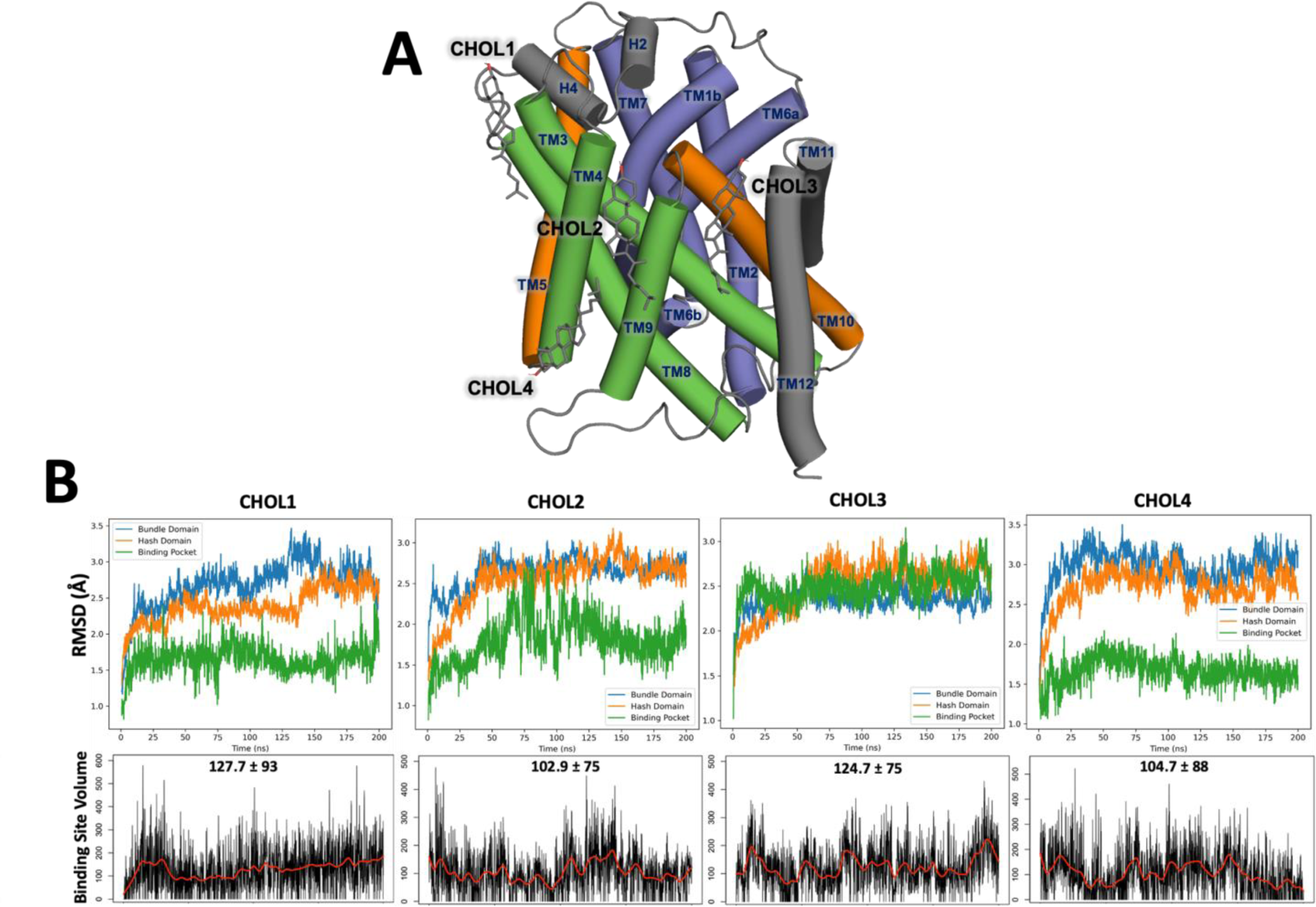
Correlated movement of LAT1’s functional motifs and substrate binding site with cholesterol binding. **(A)** Structural overview of cholesterols relative to LAT1 functional motifs. The bundle domain (blue) of LAT1 comprises of TMs 1, 2, 6, 7, the hash domain (green) TMs 3, 4 8, 9 and the arms (orange) TMs 5,9. LAT1 functional motifs are displayed as cartoon, while the cholesterols are represented as sticks. **(B)** Comparison of the effect of each cholesterol molecule on two functional motifs of LAT1: The stability of the residues (green) in the substrate binding pocket and the evolution (below) of the volume of that pocket, is evaluated during the simulation. The average volume of the substrate binding site of each simulation is added in bold. The x-axis displays the duration of the simulation trajectory (in nanoseconds), and the y-axis conveys the movement of atoms in angstroms (Å), compared to the initial frame of the trajectory (above) and the size of the binding pocket (below) in angstroms (Å).

Interestingly, CHOL3 is located near the hash domain (TM3 and TM9), which may explain the increased stability and coordinated movement of the motifs and substrate binding site. Additionally, we observe CHOL3 maintains a larger orthosteric substrate binding site (124.7 ± 75 Å^3^) during the simulation, when compared to CHOL2 and CHOL4 (102 ± 75 Å^3^ and 104.7 ± 88 Å^3^, respectively).

Interestingly, during the CHOL1 simulation, the orthosteric substrate binding site is similar in volume to the corresponding site in the CHOL3 simulation (127.7 ± 93 Å^3^ and 124.7 ± 75 Å^3^, respectively), however, the motifs and substrate binding site appear to be less stable in the CHOL1 simulation than in the CHOL3 simulation (Fig. 3). This difference is likely due to time it takes for cholesterol to settle into the slightly lower binding pocket as discussed previously. Of note, neither the CHOL2 or CHOL4 simulation showed increased stability or synchronization of the motifs and binding site residues observed in the CHOL3 simulation.

### Docking and free energy of binding of cholesterol to CHOL1-4

To elucidate the binding affinity of cholesterol to the identified sites within LAT1, we performed molecular docking and binding free energy calculations. First, we docked a cholesterol molecule to each of the CHOL1-4 sites. CHOL3 site yielded the most favorable docking score of -5.55, which not only demonstrated the best fit but also showed a deeper cholesterol binding pose within the binding pocket, as evidenced by a RMSD of 2.9 Å from the cryo-EM structure. This observation aligns favorably with MD simulation results and the structural dimensions of the CHOL3 site, which suggest its capability to accommodate cholesterol effectively (Suppl. Fig 3C).

Conversely, CHOL4 and CHOL2 exhibited suboptimal docking scores of -1.94 and -3.92, respectively. The docking poses were also mismatched to the observed pose in the cryo-EM structure (CHOL2 RMSD = 2.7 Å; CHOL4 RMSD = 1.6 Å), with CHOL2 presenting a nearly perpendicular orientation and CHOL4 exhibiting a flipped cholesterol head group.

Interestingly, docking to CHOL1 produced an intriguing result, where the docking produced an unfavorable score of -2.33 despite the pose closely resembled that in the pdb structure (RMSD of 1.0 Å). This was unexpected as our previous findings suggested that CHOL1 provides sustained contacts with cholesterol and multiple residues (Figures 2,3). Prompted by this discrepancy, we explored the possibility of an alternative binding mode at CHOL1. We defined a new site called CHOL1^refined^, derived from the most prevalent cluster observed during the CHOL1 MD simulations. Remarkably, this refinement led to a significantly improved docking score of -5.30, with an excellent alignment to the CHOL1^refined^ structure(RMSD = 1.0 Å).

To complement the docking studies, we estimated the binding free energies using the MM/GBSA method for each docking pose. Consistent with the docking scores, CHOL3 exhibited the lowest free energy of binding at -49.96 kcal mol^-1^, suggesting the strongest interaction with cholesterol. Comparatively, CHOL1, CHOL2, and CHOL4, showed higher binding free energies, indicating weaker interactions. Notably, CHOL1refined achieved the lowest binding free energy of -54.30 kcal mol^-1^.

These results collectively indicate that both CHOL3 and the revised CHOL1^refined^, offer the highest complementarity and potentially the strongest binding affinities for cholesterol, highlighting their significance in modulating LAT1 function.

## DISCUSSION

Describing the structural determinants for LAT1-cholesterol recognition is crucial for understanding cholesterol’s role in modulating LAT1 function and provide a framework for the development of novel small molecule modulators of LAT1. This study deployed computational methodologies to define the putative cholesterol binding sites, elucidate LAT1-cholesterol interactions, and illustrate the influence of cholesterol on the structural dynamics of LAT1.

First we assessed whether CHOL1-4 ^33^ are complementary to cholesterol in terms of structural features such as size and shape. Our results point to CHOL3 as the premier candidate for cholesterol accommodation due to its superior surface area, volume, and hydrophobic characteristics, which are conducive to fit snugly into the pocket (Fig. 1B, Suppl. Table 1). MD Simulations further support this, demonstrating a stable and deep engagement of cholesterol, binding deeper into this large groove than observed in the experimentally determined cryo-EM structure (Suppl. Fig 3). Additionally, the MD simulations underscore the stability of cholesterol at CHOL3, where it maintains persistent and specific interactions with coordinating residues across three cholesterol moieties for over 90% of the simulation duration (Fig. 3). CHOL1, while exhibiting a similarly sized groove, appears to favor cholesterol binding at CHOL1^refined^ conformation rather than the published structure.

Next, we explored whether cholesterol’s binding to LAT1 correlated with the dynamics of LAT1’s functional motifs (i.e., bundle and the hash) and the substrate binding site. Previous studies^14–18, 33^ have shown cholesterol plays a role in modulating the function of related transporters such as SERT and DAT, suggesting a similar phenomenon may occur for LAT1. For example, Laursen et. al. elucidated that direct cholesterol binding to SERT, shifted the transporter conformational equilibrium to the outward open state^17^. Another study by Zeppelin et. al., focusing on the dopamine transporter, proposed cholesterol binding inhibited the outward-to-inward transition. For LAT1, the rocking bundle, motions of TM1 and TM6, are essential for conformational transitions between the inward open and outward open states. This is defined by a rotation of these TMs to swing in or out, thus exposing the substrate site of the protein to either the intracellular or extracellular side of the cell^33, 35, 52^.

Our simulations indicate that cholesterol binding at CHOL3 appears to stabilize the functional motifs and substrate binding site, suggesting coordinated movement (Fig. 4). It is plausible that the increased stability, may contribute to the observed ability of cholesterol to increase the Vmax and not the Km of LAT1 mediated uptake of substrates^19^. Taken together, we propose that CHOL3 and to a lesser extent CHOL1^refined^ are functionally important.

In recent years, LAT1 has emerged as a viable drug target for multiple diseases and disorders such as cancer^53–56^ and autism spectrum disorder (ASD)^57, 58^. Currently, most of drug discovery efforts targeting LAT1 has focused on inhibitors targeting the substrate binding site. For example, there are two substrate-binding site inhibitors of LAT1 that are currently under clinical investigation for treating patients with advanced biliary tract cancers (JPH-203)^59^ and advanced pancreatic cancer (OKY 034)^60^. One limitation of targeting the substrate binding sites of transporters is their limited selectivity compared to transporters with similar substrate profiles as well as the lack of drug-like properties of amino-acid like compounds^61^. Therefore, potential allosteric binding sites (ex: non-competitive inhibition) can be beneficial, due to the specificity and lower conservation an allosteric site may provide over the substrate site and inhibition of function regardless of orthosteric substrate concentration. Recent studies have shown other SLC transporters (e.g., SLC1 family^62^) can be targeted with an allosteric binder to modulate protein function.

A prerequisite for the development of future chemical tools and drugs targeting a LAT1 allosteric site, is to characterize its druggability, i.e. their ability to bind small-molecule compounds. To begin to answer this question, we evaluated the docking affinity and binding free energy of cholesterol across these sites. These results revealed that to no surprise, CHOL3 provided superior docking and free energy of binding profiles (Fig. 5). Meanwhile, CHOL1 initially showed underwhelming results but CHOL1^refined^ displayed a remarkable enhancement in both docking affinity and binding free energy.

**Figure 5:**
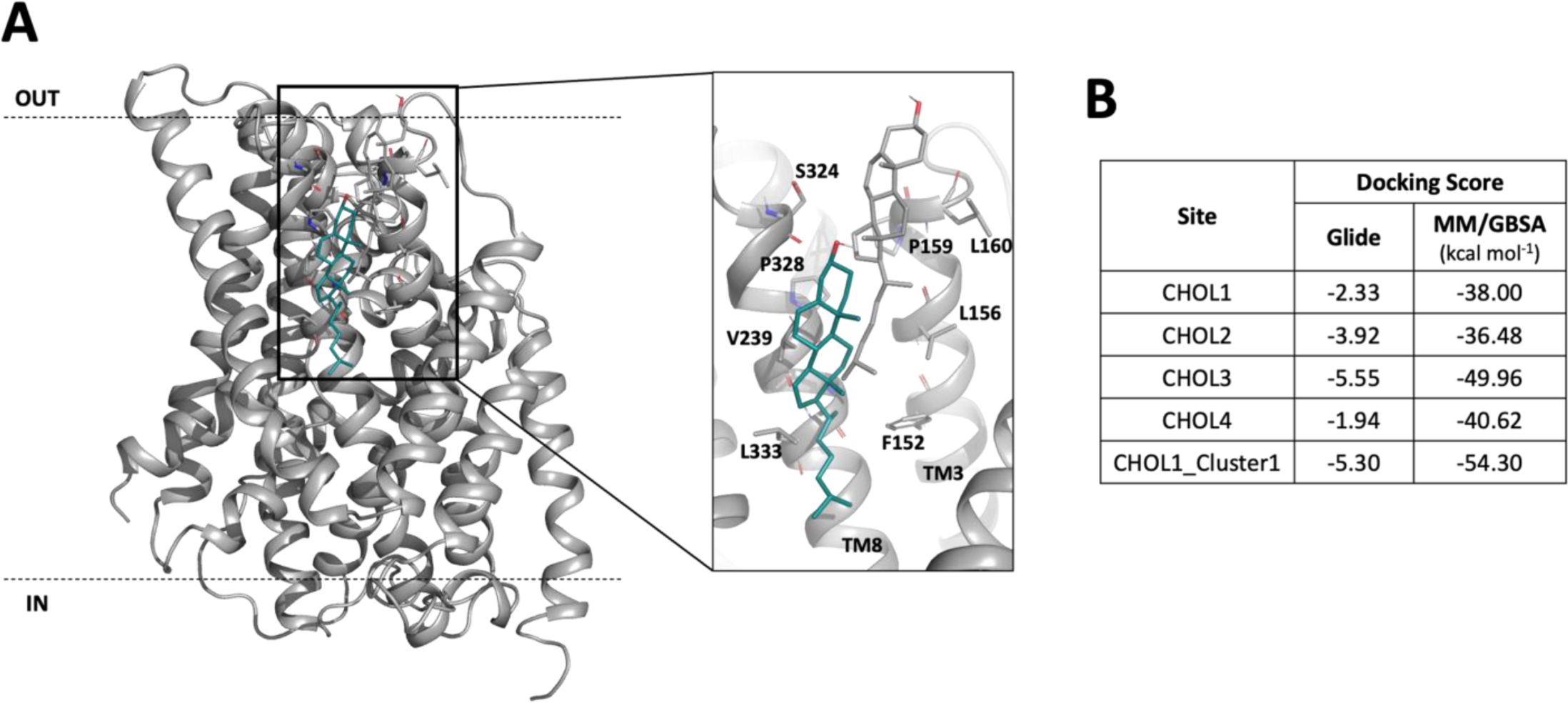
Predicted LAT1-cholesterol binding using docking and free energy estimation. (**A**) Comparison of the native binding pose of CHOL1 (grey) from the cryo-EM structure, and the representative binding pose (teal) of cholesterol from the dominant cluster of the CHOL1 MD simulation shown in Fig. 2. (**B**) Docking score obtained via Glide and the free energy of binding obtained from Molecular Mechanics with generalized Born and surface area solvation (MM/GBSA), for each cholesterol site (CHOL1-4). The docking and MM/GBSA scoring of cholesterol docked to the modified CHOL1 binding site based on the MD simulation is also described.

In summary our results endorse CHOL3 site as the most favorable site for cholesterol binding, with CHOL1^refined^ also demonstrating significant potential. We propose both these sites can be targeted for small molecule discovery, which may allow for further unravelling of the regulatory role of cholesterol in LAT1’s function and modulation.

