## Supplementary Figures 1-4 for "Mechanisms of Cholesterol Binding to LAT1"

**SUPPLEMENTARY FIGURE 1**

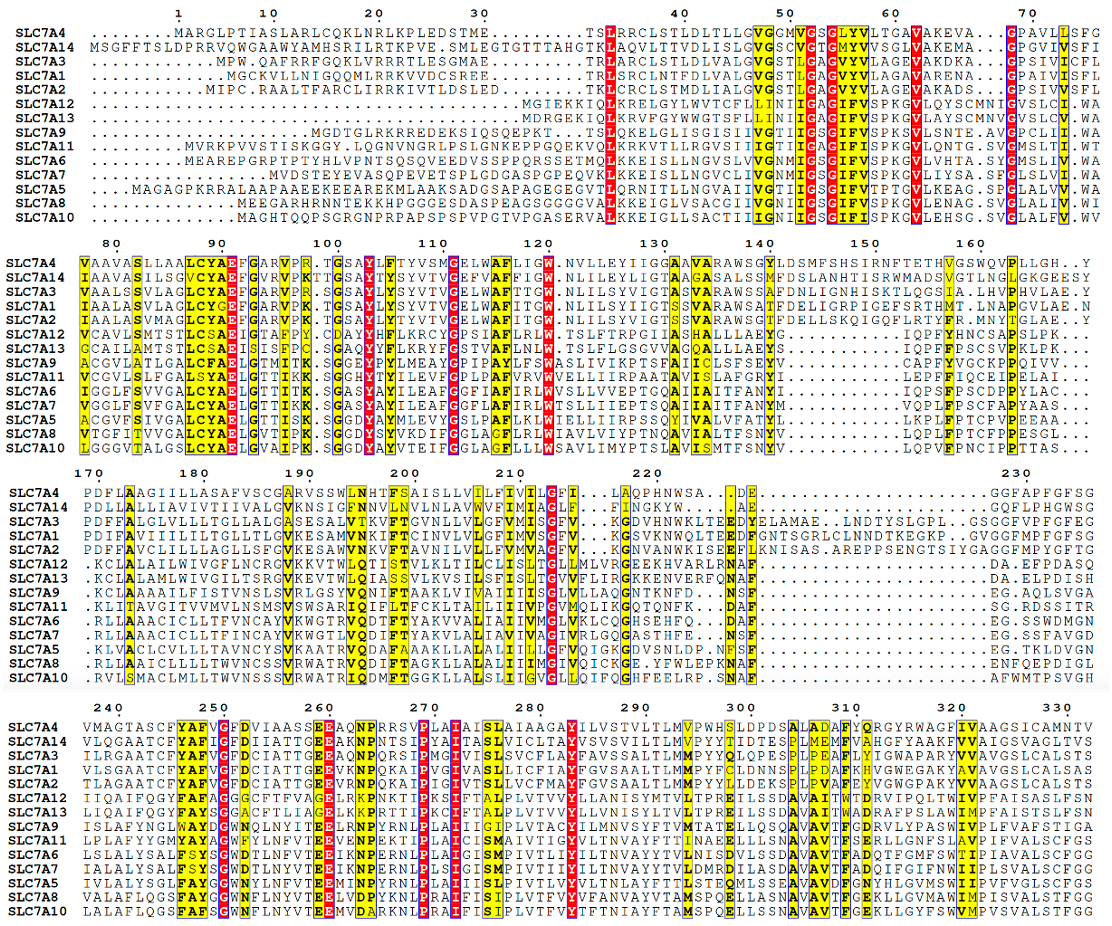

**Supplementary Figure 1: MSA of SLC7 Family.** Multiple sequence alignment of each member of the SLC7 family generated by MUSCLE. Residues that have 100% sequence identity are highlighted in red, whereas residues with high sequence similarity are highlighted in yellow.

**SUPPLEMENTARY FIGURE 2**

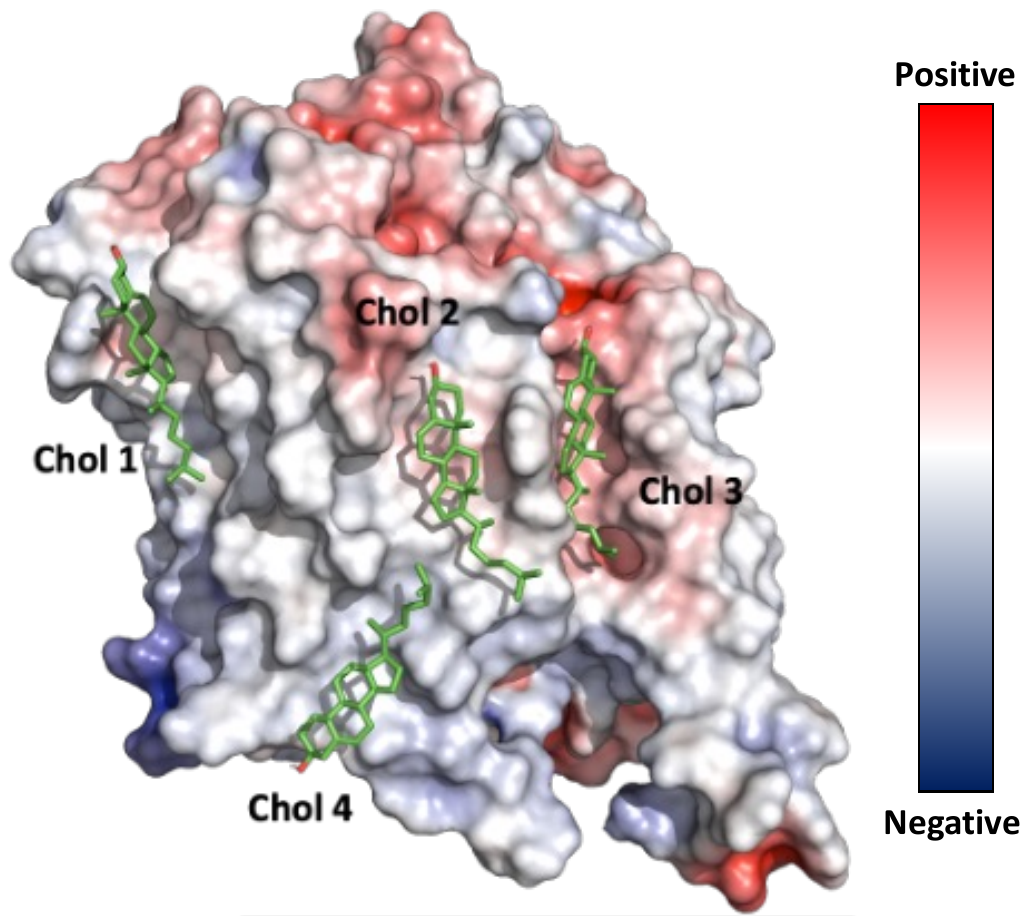

**Supplementary Figure 2:** **CHOL1-4 electrostatic profile.** Electrostatics analysis of the cryo-EM structure of LAT1 in the apo inward-open conformation (PDB: 6JMQ). Each putative cholesterol is represented as sticks, with the LAT1 protein in surface representation.

**SUPPLEMENTARY FIGURE 3**

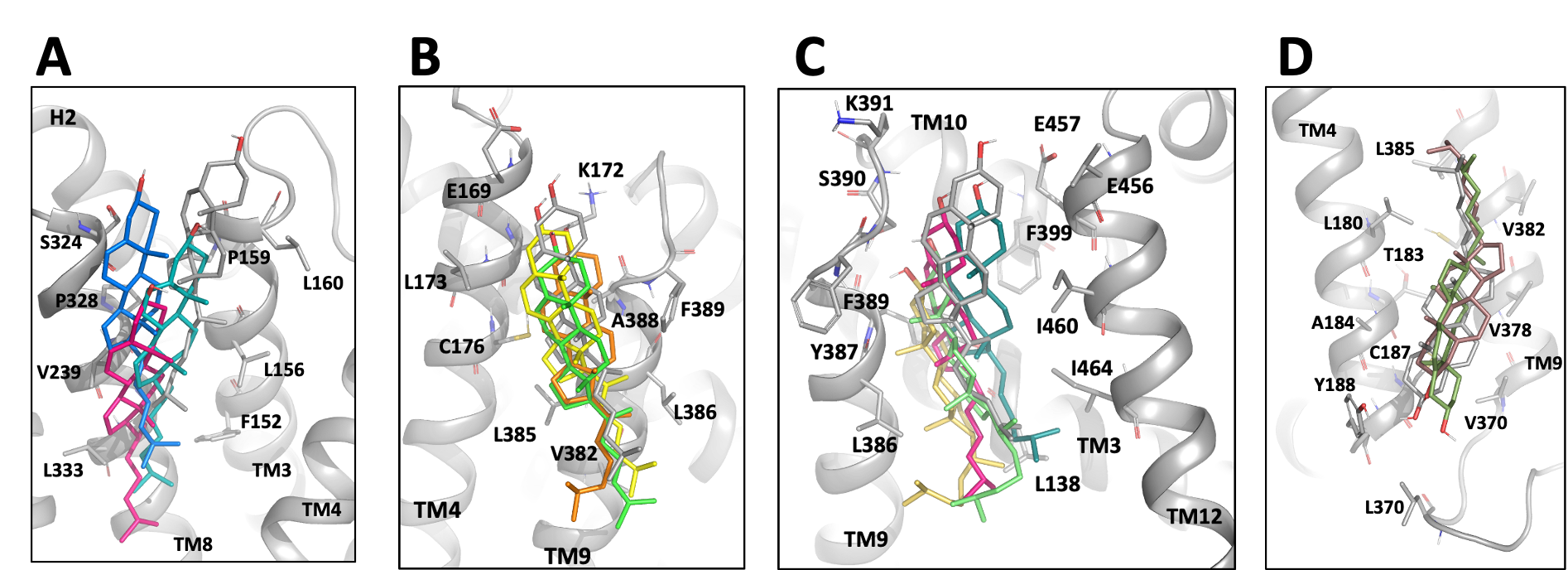

**Supplementary Figure 3: Cluster analysis of CHOL1-4 MD Simulations.** Ensemble conformations of LAT1 generated from CHOL1-4 (**A-D**) simulation are shown. Superposition of the apo inward-open conformation structure (PDB:6JQM) in grey and each labeled individual representative model from each cluster (colored).

**SUPPLEMENTARY TABLE 1**

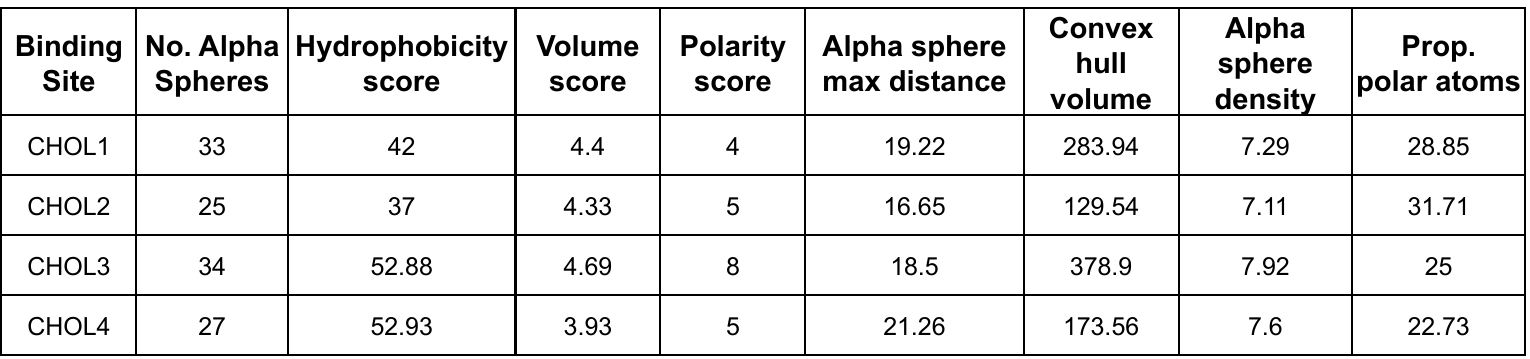

**Table 1:** Table showing generated molecular descriptors of CHOL1-4 sites, calculated by dpocket.
